# The Sec61 translocon is a therapeutic vulnerability in Multiple Myeloma

**DOI:** 10.1101/2021.09.05.459026

**Authors:** Antoine Domenger, Caroline Choisy, Ludivine Baron, Véronique Mayau, Ludovic Deriano, Bertrand Arnulf, Jean-Christophe Bories, Gilles Dadaglio, Caroline Demangel

**Affiliations:** Unité ‘Immunobiologie de I’Infection’, Institut Pasteur, INSERM U1221, 75015 Paris, France; Université de Paris, Sorbonne Paris Cité, 75013 Paris, France; INSERM U976 Équipe 5, Institut de Recherche Saint Louis, Université de Paris, 75010 Paris, France; Unité ‘Intégrité du Génome, Immunité et Cancer’, Equipe Labellisée Ligue Contre Le Cancer, Institut Pasteur, INSERM U1223, 75015 Paris, France; APHP Department of Immuno-Hematology, Hôpital Saint Louis, 75010 Paris, France

**Keywords:** Multiple Myeloma, Sec61 translocon, Proteostatic stress

## Abstract

Multiple Myeloma (MM) is an incurable malignancy characterized by the uncontrolled expansion of plasma cells in the bone marrow. While proteasome inhibitors like bortezomib efficiently halt MM progression, drug resistance or toxicity inevitably develop. Here, we used a recently discovered inhibitor, mycolactone, to assess the interest of targeting the Sec61 translocon in MM. In human cell lines and tumors from MM patients, mycolactone triggered pro-apoptotic endoplasmic reticulum stress responses synergizing with bortezomib for induction of MM cell death, irrespective of their resistance to proteasome inhibition. Notably, this synergy was selective of cancer cells and extended to B cell acute lymphoblastic leukemia. Sec61 blockade also caused collateral defects in MM secretion of immunoglobulins and expression of pro-survival interleukin (IL)-6 receptor and CD40, whose activation stimulates IL-6 production. Further, the mycolactone-bortezomib combination demonstrated superior therapeutic efficacy over single drug treatments in immunodeficient mice engrafted with MM cells, without inducing toxic side effects. Collectively, these findings establish Sec61 blockers as novel anti-MM agents and reveal the interest of targeting both the translocon and the proteasome in proteostasis-addicted tumors.

## Introduction

Multiple Myeloma (MM) is a malignant disorder characterized by the uncontrolled expansion of clonal plasma cells in the bone marrow, eventually leading to organ dysfunction and death (1). The global burden of MM has increased by 126% over the past 30 years, with highest incidences in developed countries (2). The introduction of proteasome inhibitors (PIs) like bortezomib (BZ) in MM chemotherapies has revolutionized MM management, and combinations of BZ with immunomodulators and dexamethasone have become the gold standard for MM treatment (3). However, BZ can induce severe side effects such as peripheral neuropathy requiring discontinuation of therapy, and many patients develop resistance to BZ whose molecular basis remains poorly understood (4). Most importantly, treated patients relapse and less than half of them survive beyond 5 years post-diagnosis. Second-generation of PIs were recently developed to treat patients that are resistant or intolerant to BZ (5), but these drugs induce other types of side effects and do not prevent relapse. Therefore, despite significant therapeutic advances, MM remains an incurable disease and the identification of new therapeutic targets is critically needed.

The clinical efficacy of BZ is primarily attributed to its ability to induce the accumulation of misfolded immunoglobulins in the cytoplasm of MM cells, leading to lethal proteotoxic stress (5). BZ also alters the survival and proliferation of MM cells in several other ways, such as inhibition of NF-κB oncogenic signaling, suppression of pro-adhesive cross-talks with bone marrow stromal cells and prevention of growth stimulation by cytokines like Interleukin (IL)-6 (6, 7). Amongst the different strategies that are currently explored to complement or replace PIs in MM chemotherapies are novel means to disrupt the homeostatic regulation of the secretory apparatus, via proteolytic routes and endoplasmic-reticulum (ER)-associated protein degradation (ERAD) components other than the proteasome (8). Despite its central importance in MM biology, the interest of targeting immunoglobulin transport into the ER has not yet been explored, due to the lack of suitable inhibitor.

We reported recently that mycolactone, a diffusible lipid toxin secreted by the human pathogen *Mycobacterium ulcerans*, operates by inhibiting the mammalian translocon (Sec61) (9, 10). By targeting the central pore-forming subunit of Sec61 (Sec61α), mycolactone prevents the import of newly synthesized secreted and transmembrane proteins into the ER, leading to their cytosolic degradation by the proteasome (11–13). Contrary to the Sec61 inhibitor cotransin (14), mycolactone is not substrate-selective and blocks the translocation of the vast majority of Sec61 clients (9, 15). The only substrates resisting its inhibitory action are transmembrane proteins belonging to the rare Type III subgroup (10-12, 15). Within one hour of treatment, mycolactone-treated cells become defective for production of most secreted proteins and membrane-anchored receptors. If sustained, mycolactone treatment triggers proteotoxic stress responses in cytosol and ER, ultimately inducing apoptosis (15, 16). Notably, a point mutation (R66G) in the Sec61α amino acid sequences preventing mycolactone binding without affecting the translocon functionality was sufficient to prevent stress responses and associated cytotoxicity, demonstrating that Sec61 inhibition by mycolactone is the molecular mechanism driving cell death (9).

Based on these findings, we hypothesized that Sec61 blockade may represent a novel targeted therapy in MM, suppressing the survival and growth of MM cells in at least two ways: by generating lethal proteotoxic stress and by preventing the expression of membrane receptors that are key to MM cell division and dissemination. The present study uses mycolactone as a prototypical Sec61 blocker to establish proof-of-concept, and evaluate the translational potential of Sec61 inhibitors in the treatment of MM.

## Results

### Sec61 blockade by mycolactone severely alters the biology and viability of MM cell lines

To assess the effect of mycolactone on MM cell viability, three human cell lines (MM.1S, JIM3 and KMS-11) were treated with increasing concentrations of mycolactone for 24-72 h, and the induction of cell apoptosis was monitored by phosphatidylserine exposure (Annexin V staining) and loss of membrane integrity (PI staining) (**Fig. S1**). While the three cell lines differed in sensitivity to mycolactone, MM.1S being the most resistant, a dose- and time-dependent induction of apoptotic cell death was consistently observed after 48 h of treatment (**Fig. 1A**). In all cell lines, initiation of apoptosis was preceded by a decrease in surface expression of the MM cell marker CD38 (17) (**Fig. 1B**). The plasma cell marker CD138 supports MM cell survival in the bone marrow by promoting growth factor signaling (18). While not detected on JIM3 and KMS-11, CD138 expression by MM.1S was also dose-dependently decreased by mycolactone after 24 h (**Fig. 1B**). IL-6 receptor (IL-6R) and CD40 are two other MM cell markers whose signaling play a crucial role in MM development and dissemination (7, 19). When expressed by the MM cell lines, these receptors were also rapidly and potently down-regulated by mycolactone treatment (**Fig. 1B**).

**Figure 1:**
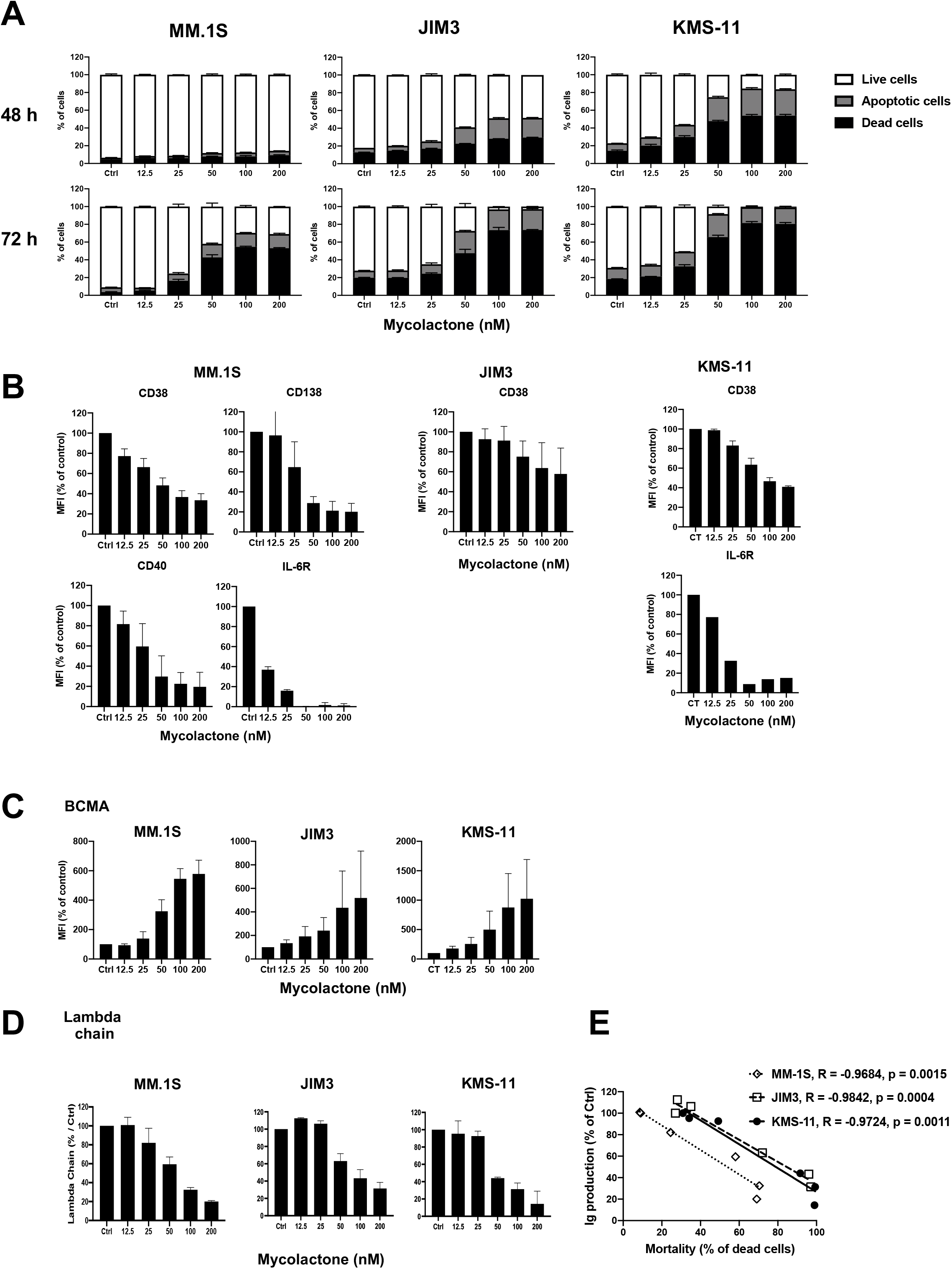
Sec61 blockade by mycolactone strongly alters the biology and viability of MM cell lines. **(A)** Effect of mycolactone on MM.1S, JIM3 and KMS-11 cell viability after 48 and 72 h of treatment, as measured by exposure of Annexin V and incorporation of PI. Live, apoptotic and dead cells were determined as in Fig. S1. Data are Means ± SD of duplicates, expressed as percentages of total cells. **(B)** Inhibitory effect of mycolactone on cell expression of CD38 (Type II) and CD138, IL-6R, and CD40 (Type I) transmembrane proteins. **(C)** Stimulatory effect of mycolactone on cell expression of BCMA, a Type III transmembrane protein. Mean Fluorescence Intensities (MFIs) in **(B)** and **(C)** were measured in live cells 24 h after addition of mycolactone. Data are Mean MFIs ± SD from 2 independent experiments, relative to vehicle-treated controls (Ctrl). **(D)** Inhibitory effect of mycolactone on immunoglobulin lambda chain secretion, as quantified by ELISA of culture supernatants after 24 h of treatment. Data are Mean ± SD of duplicates of secreted lambda chain (μg.mL^-1^) concentrations, relative to vehicle-treated controls (Ctrl). **(E)** Correlation between immunoglobulin lambda chain (Ig) secretion after 24 h (D) and MM cell line mortality after 72 h (A), in the 3 cell lines treated with 50 nM mycolactone. Slopes (R) and statistical significance (p values) are indicated. All panels in Figure 1 are representative of at least 2 independent experiments with similar results.

In contrast to CD38, CD138, IL-6R and CD40, all Type I or II transmembrane proteins efficiently blocked in translocation by mycolactone, Type III transmembrane proteins are Sec61 substrates resisting mycolactone inhibition (9, 11, 12, 15). B-cell maturation antigen (BCMA) is a Type III protein that is typically expressed by MM cells and the target of next-generation immunotherapies (20). In all cell lines, mycolactone dose-dependently increased surface expression of BCMA (**Fig. 1C**). On the opposite, Sec61 inhibition efficiently decreased MM cell line secretion of immunoglobulin light chains after only 24 h of treatment (**Fig. 1D**), and this reduction closely correlated with the onset of cell death after 72 h (**Fig. 1E**). In conclusion, Sec61 blockade by mycolactone induces programmed cell death in MM preceded by phenotypic defects in expression of Type I/II membrane receptors and secretion of immunoglobulins.

### Mycolactone synergizes with proteasome inhibitors for induction of MM cell line apoptosis

The clinical efficacy of BZ is believed to result from its capacity to induce unresolvable proteotoxic stress in MM cells via accumulation of misfolded proteins, including immunoglobulins, in the cytoplasm (5). Since Sec61 substrates that are blocked in translocation by mycolactone are diverted to the proteasome for cytosolic degradation (13), we reasoned that mycolactone may potentiate the anti-MM activity of BZ cells through generation of additional proteotoxic stress. As shown in **Figure 2A**, the three MM cell lines were differentially susceptible to BZ treatment, JIM3 and KMS-11 being relatively more resistant than MM.1S. To assess a potential synergy between the two drugs, MM cell lines were treated for 24 h with sub-toxic doses of BZ together with increasing concentrations of mycolactone. In all cell lines, the mycolactone-BZ combination induced more cell death than single drugs (**Fig. 2B and S2**). Heatmaps of synergy scores (21) show that mycolactone synergized with BZ in all cell lines, irrespective of their basal resistance to BZ (**Fig. 2C**). Similar findings were obtained with MM.1S cells treated with carfilzomib, a second-generation PI that is proposed to BZ-resistant MM patients (**Fig. 2D, 2E**).

**Figure 2:**
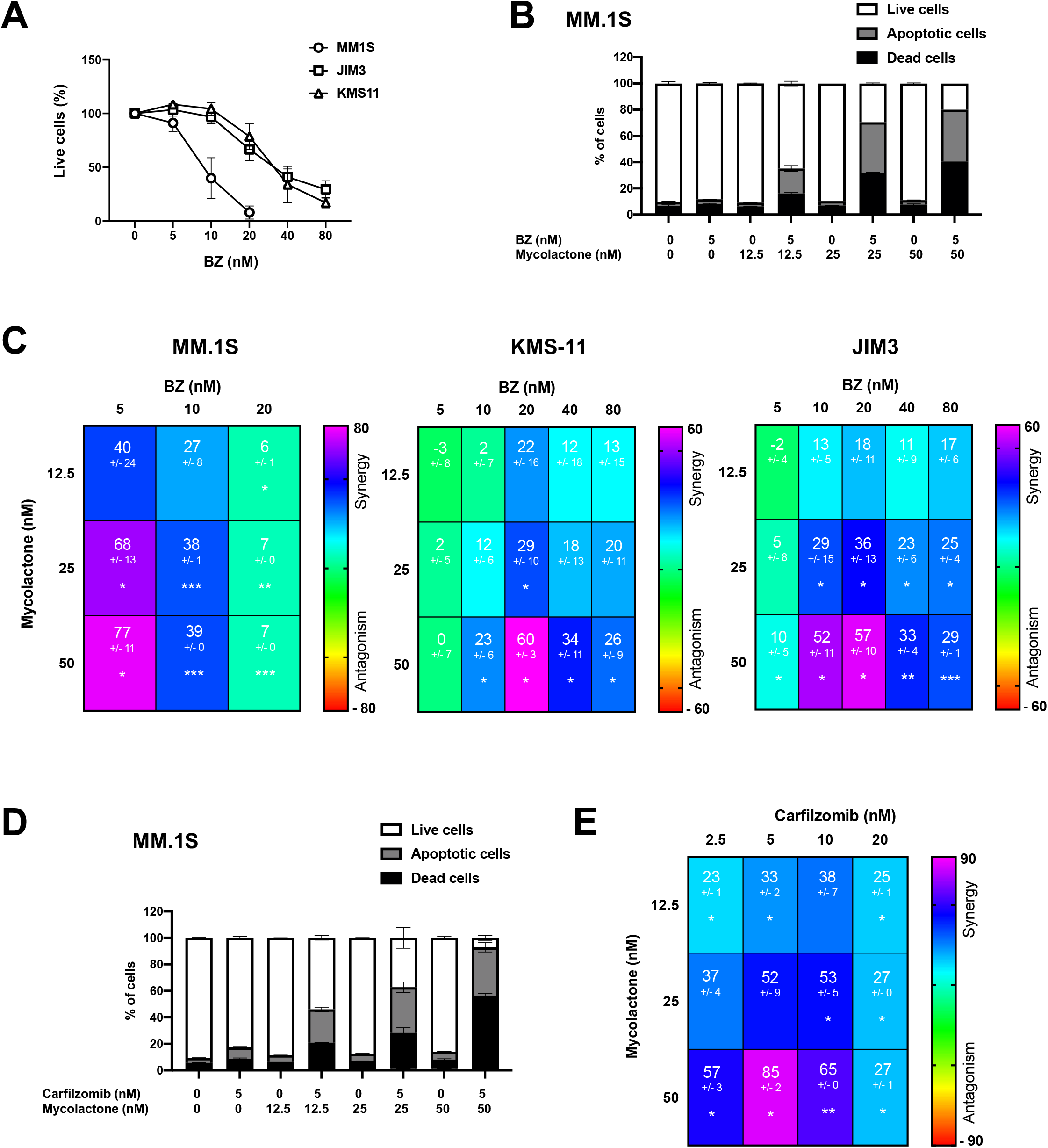
Mycolactone synergizes with PIs for induction of MM cell apoptosis. **(A)** MM.1S, JIM3 and KMS-11 cells were incubated with increasing concentrations of BZ for 24 h. The proportion of live (Annexin V^-^ PI^-^) cells was then measured by flow cytometry as in Fig. S1. Data are Mean % of live cells +/- SD of duplicates, relative to vehicle-treated controls (Ctrl), and are representative of 2 independent experiments with similar results. **(B)** MM.1S were treated for 24 h with mycolactone and/or BZ at the indicated concentrations. Percentages of live, apoptotic and dead cells were then measured by flow cytometry as in Fig. S1. Data are Mean % ± SD of duplicates, relative to total cell number, and are representative of 3 experiments with similar results. **(C)** Synergy between mycolactone and BZ in the MM.1S, JIM3 and KMS-11 cell lines, when treated as in (B). Data are Mean Loewe scores ± SD, shown as heatmaps. Differences between treated cells and controls by Student’s t-test: *\*p<0.05; **p<0.01; ***p<0.001*, N = 6 (cumulative data of 3 independent experiments with two duplicates) for MM.1S and KMS-11 and N = 4 (cumulative data of 2 independent experiments with two duplicates) for JIM3. **(D and E)** MM.1S cell were treated as in (B) but using carfilzomib instead of BZ. Synergy scores were calculated as in (C), N = 4 (cumulative data of 2 independent experiments with two duplicates).

### Combining mycolactone with BZ enhances ER stress in MM cell lines

We next sought to determine if the synergistic induction of apoptosis in MM cell lines treated with mycolactone and BZ was associated with enhanced ER stress. The Unfolded Protein Response (UPR) is activated by three ER-resident transmembrane proteins: Inositol-Requiring Enzyme 1 (IRE1), Protein Kinase RNA (PKR)-like ER Kinase (PERK) and Activating Transcription Factor (ATF) 6 (**Fig. 3A**). Activation of IRE1 triggers the splicing of X-box binding protein 1 (XBP1) mRNA into its transcriptionally active form sXBP1; while that of PERK up-regulates ATF4, and ATF6 is processed into an active form in the Golgi. ER stress-driven apoptosis is largely mediated by induction of C/EBP homology protein (CHOP) gene expression, activated by sXBP1, ATF4 and ATF6. Protein levels of ATF4 and mRNA levels of CHOP and sXBP1 were all potently induced by the mycolactone-BZ combination after only 6 h of treatment, while single drugs had minor effects at this time point (**Fig. 3B, 3C**). The sharp induction of UPR markers after 6 h correlated with the onset of cell death observed 18 h later (**Fig. 2**). Together, these data suggested that enhanced ER stress contributes to the cytotoxic synergy between mycolactone and BZ.

**Figure 3:**
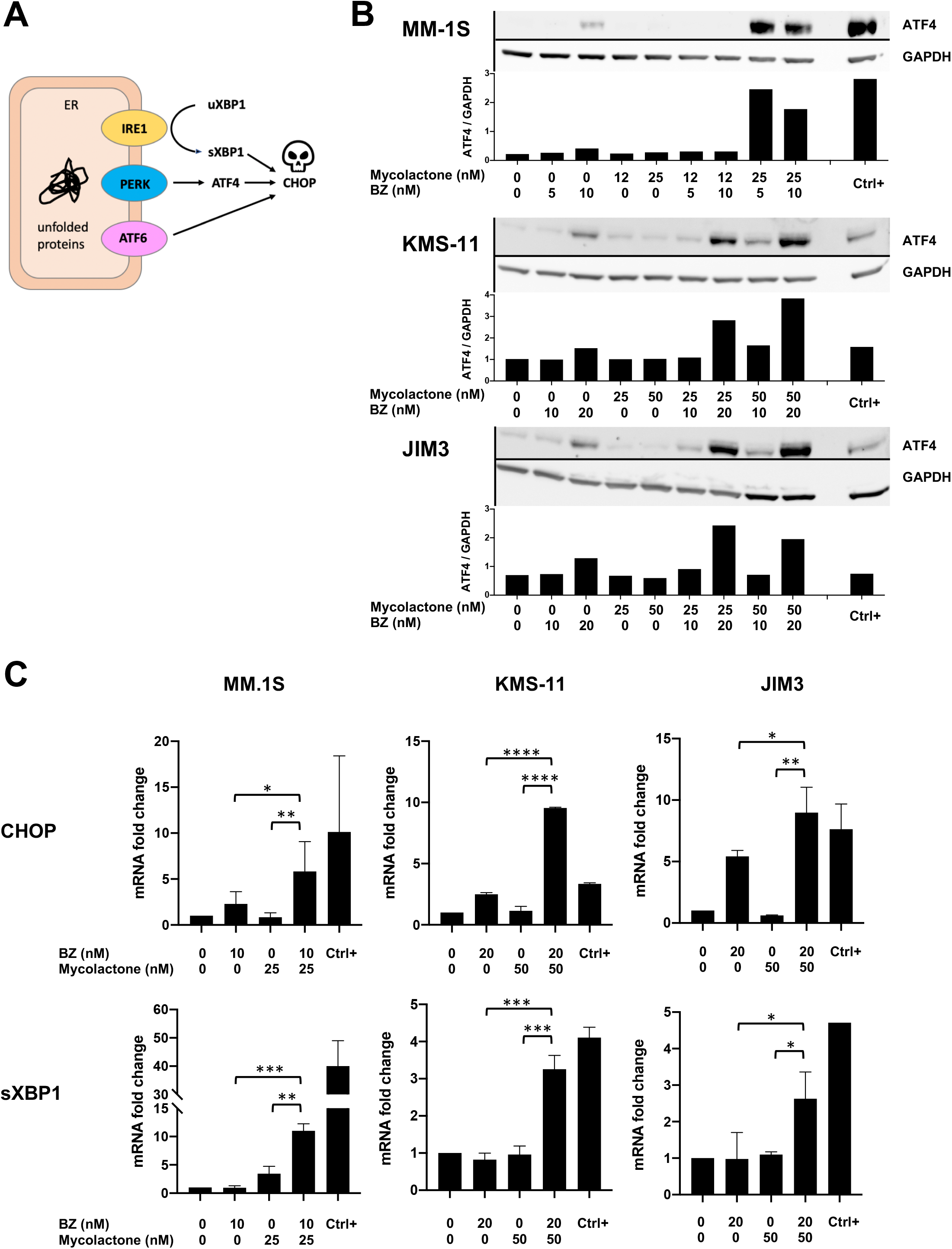
Combining mycolactone with BZ enhances ER stress in MM cell lines. **(A)** Diagram illustrating the pro-apoptotic pathways activated by UPR sensors IRE1, PERK and ATF6 upon accumulation of unfolded proteins in the ER lumen. **(B)** MM.1S, KMS-11 and JIM3 cells were treated with mycolactone and/or BZ, the ER stress inducer Thapsigargin (2 μM) as positive control (Ctrl^+^) or vehicle as negative control, for 6 h. ATF4 protein levels were assessed in cell lysates by Western blot (top panel) and quantified relatively to GADPH levels (lower panel). Data are representative of two independent experiments with similar results. **(C)** CHOP and sXBP1 mRNA levels were quantified by qPCR in the three cell lines treated as in (B). sXBP1 mRNA levels were normalized to total (spliced + unspliced) XBP1 mRNA level. Data are Mean RNA fold changes (2^-ΔΔCT^) ± SD relative to untreated controls, calculated from two independent experiments and pairwise compared by one-Way ANOVA with Fisher’s LSD. *\*p<0.05; **p<0.01; ***p<0.001, ****p<0.0001*.

### The synergy between mycolactone and BZ cytotoxicity extends to mouse B cell lymphomas

Besides MM, PIs show promise in the treatment of other hematological malignancies such as acute leukemia (22), and solid malignancies (23). We tested the proteotoxic effects of mycolactone, alone and combined with BZ, in B cell acute lymphoblastic leukemia (B-ALL) using mouse pro-B cell lines generated by transformation of hematopoietic cells with the murine viral form of the Abelson oncogene (v-abl). Mycolactone alone demonstrated potent ability to induce v-abl cell apoptosis *in vitro* (**Fig. 4A**). Moreover, as in human MM cell lines, it synergized with BZ for induction of v-abl cell death (**Fig. 4B**) and this correlated with activation of the ER stress-associated ATF4/CHOP apoptotic pathway (**Fig. 4C, 4D**). In addition to illustrating mycolactone’s ability to efficiently block mouse Sec61, these data revealed the potential interest of Sec61 inhibitors for the treatment of B-ALL, and potentially other proteasome inhibition-susceptible malignancies.

**Figure 4:**
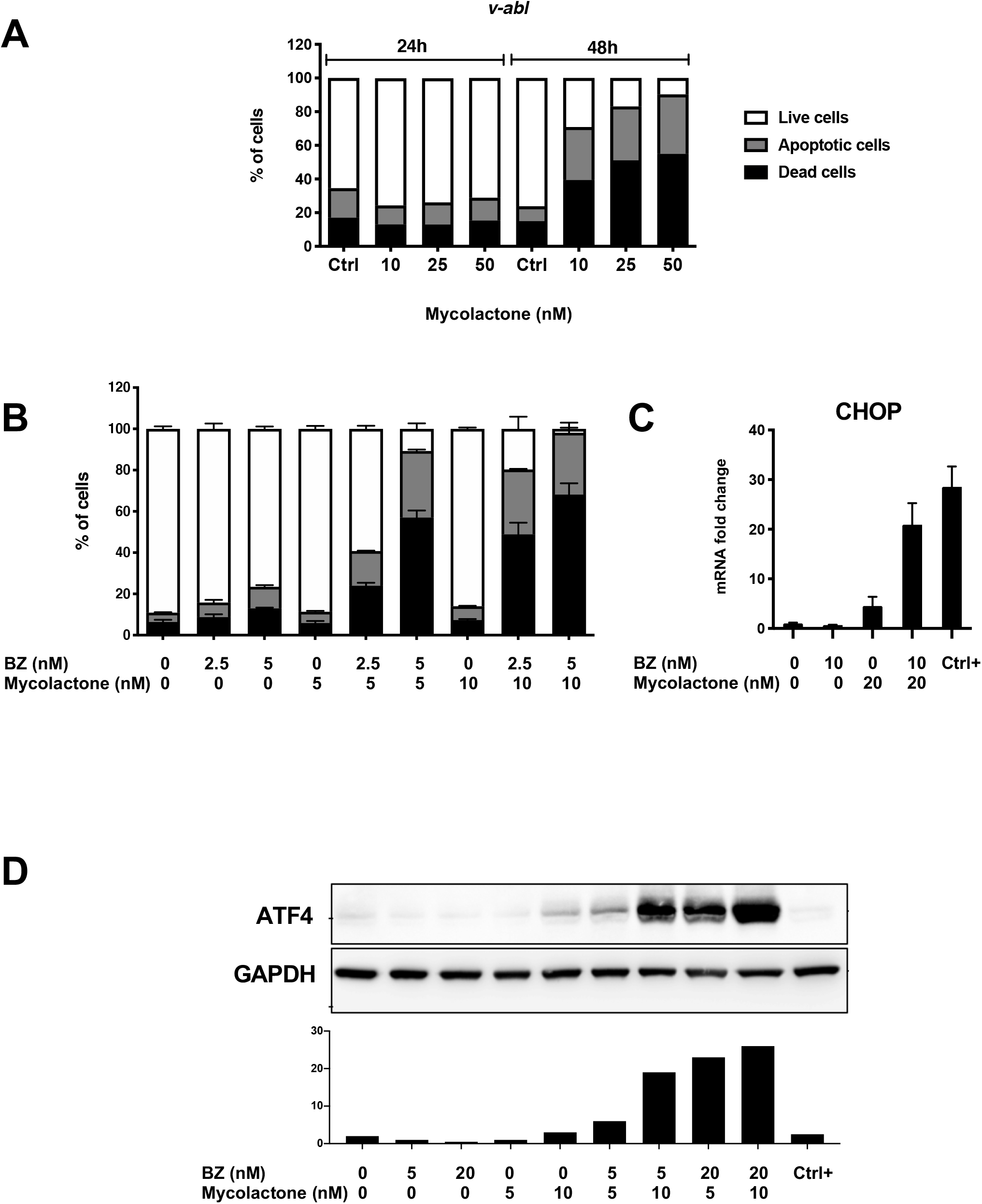
The synergy between mycolactone and BZ cytotoxicity extends to mouse B cell lymphomas. **(A)** Effect of mycolactone on v-abl pro-B cell viability after 24 and 48 h of mycolactone treatment, as measured by exposure of Annexin V and incorporation of PI. Live, apoptotic and dead cells were determined as in Fig. S1. Data are Means of duplicates, expressed as percentages of total cells. **(B)** v-abl pro-B cells were treated for 24 h with mycolactone and/or BZ at the indicated concentrations. The percentage of live, apoptotic and dead cells was then measured by flow cytometry as in Fig. S1. Data are Mean % ± SD of duplicates, relative to total cell number. **(C)** v-abl pro-B cells were treated with mycolactone and/or BZ, Thapsigargin (2 μM) as positive control (Ctrl+) or vehicle as negative control, for 6 h. CHOP mRNA levels were quantified by qPCR in the cells treated as in B. Data are Mean RNA fold changes (2^-DDCT^) ± SD relative to untreated controls, calculated from two independent experiments. **(D)** v-abl pro-B cells were treated for 6 h with mycolactone and/or BZ at the indicated concentrations. ATF4 protein levels were assessed in cell lysates by Western blot (top panel) and quantified relative to GADPH levels (lower panel). All data in Fig. 4 are representative of experiments performed with two independent v-abl cell clones with similar results.

### Patient-derived MM tumors are highly susceptible to mycolactone toxicity and synergy with BZ

We next assessed the anti-MM activity of mycolactone, alone and combined to BZ, in patient-derived tumors. Mononuclear cells were isolated from bone marrow aspirates of 4 newly diagnosed MM patients and 2 patients with relapsed MM after at least 1 line of treatment including PI and immunomodulatory drugs (IMiDs) (**Table 1**). Cells were placed in culture medium within 3 h postbiopsy, then exposed to mycolactone and/or BZ for 18 h. Induction of apoptosis was determined in MM cells, gated as CD38^+^ CD138^+/-^ plasma cells (**Fig. S3**), following Annexin V/PI staining. MM cells from the 6 studied patients varied in susceptibility to BZ treatment, irrespective of their newly diagnosed or relapsed status (**Fig. 5A and S4A**). Tumor sensitivity to mycolactone was also variable, and not systematically associated with sensitivity to BZ (**Fig. 5A and S4A**). In all studied patients, significant cell death was achieved with a 16 h exposure to ≥ 12 nM mycolactone, an anti-MM activity globally equivalent to that of 10 nM BZ. Strikingly, mycolactone synergized with BZ to kill MM cells from all patients, irrespective of their treatment-naïve or relapsed status and relative resistance to single drugs (**Fig. 5B**). Synergy was less pronounced when MM cells were highly sensitive to one of the two drugs, as in patient # 5.

**Table 1.** Characteristics of the MM patients included in this study.

| Patient | P2 | P5 | P7 | P9 | P12 | P13 |
| --- | --- | --- | --- | --- | --- | --- |
| Age (yrs) | 57 | 72 | 72 | 67 | 72 | 66 |
| Gender | F | F | F | F | M | F |
| Years since diagnosis | - | - | 10 | - | 5 | - |
| Lines of treatment | - | - | 2 | - | 1 | - |
| **Treatments** |  |  |  |  |  |  |
| IMiDs | - | - | 2 | - | 1 | - |
| PI | - | - | 2 | - | 1 | - |
| Alkylating agent | - | - | 1 | - | 1 | - |
| HSCT* | - | - | 1 | - | - | - |

\* Hematopoietic stem cell transplantation

**Figure 5:**
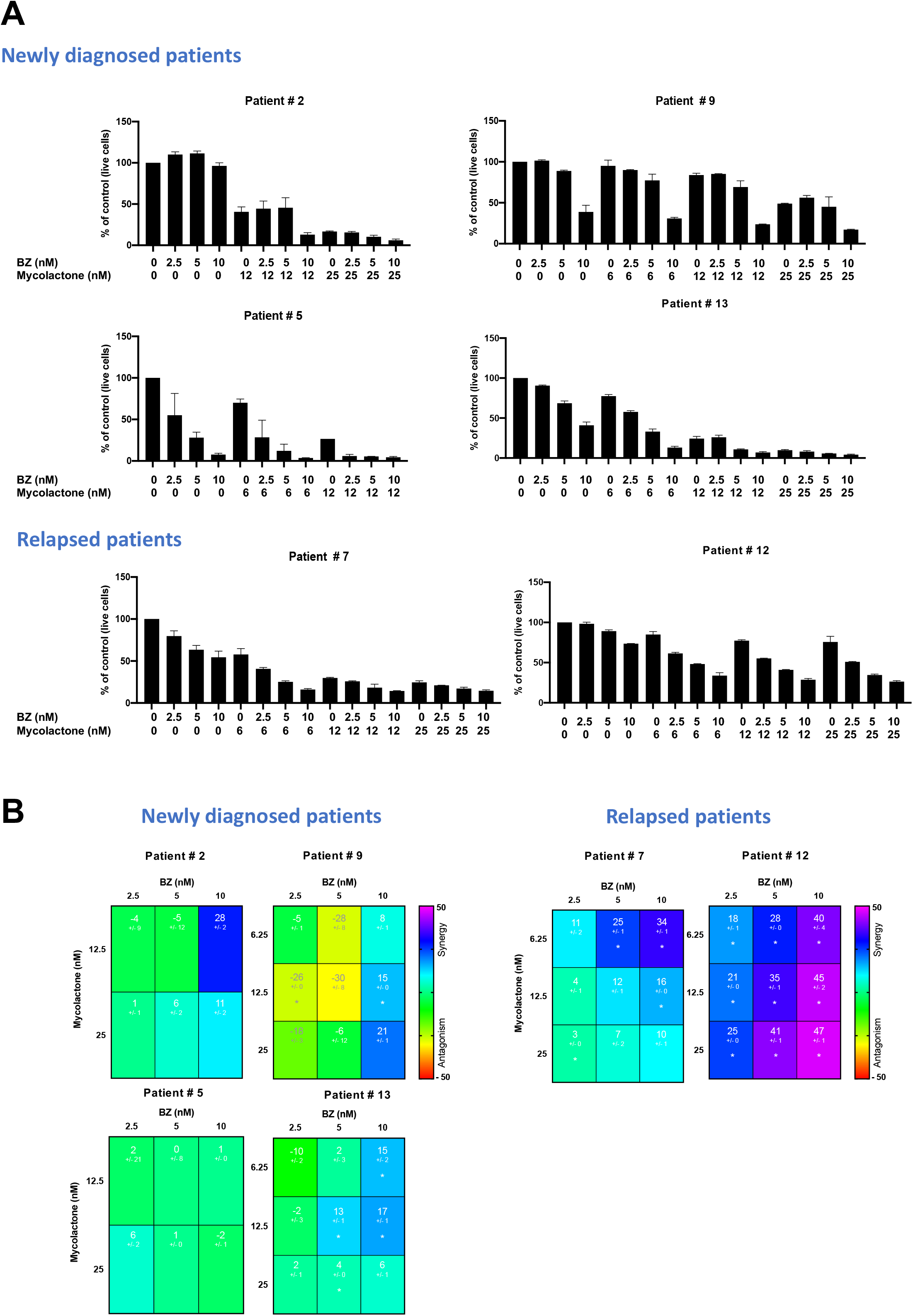
MM primary tumors are highly susceptible to mycolactone toxicity and synergy with BZ. Mononuclear cells from bone marrow aspirates of newly diagnosed or relapsed MM patients were treated with mycolactone and/or BZ at the indicated concentrations for 18 h. Then, MM cells were identified by staining with anti-CD38 and -CD138 antibodies using the gating strategy depicted in Fig. S3. **(A)** Following treatment, induction of apoptosis/necrosis was measured by exposure of annexin V and PI incorporation. Data are Means ± SD of duplicates, expressed as percentages of total cells. **(B)** Synergy between mycolactone and BZ in treated tumors. Data are Mean Loewe scores ± SD, showed as heatmaps. Differences between treated cells and controls by Student’s t-test: *\*p<0.05; **p<0.01; ***p<0.001*.

Importantly, the cytotoxicity of the mycolactone-BZ combination selectively affected MM cells, as minimal cell death was recorded in CD38^-^ CD138^-^ cells from the same bone marrow aspirates (**Fig. S4B**). It is noteworthy that bone marrow aspirates from the two relapsed patients contained a relatively higher incidence of monocytes/macrophages, as characterized by their SSC/FSC profile (as shown for Patient #7 in **Fig. S3**). Although drugs alone had little effect, the mycolactone-BZ combination displayed some toxicity on this cell subset at the highest tested concentrations (**Fig. S5A**). To further assess the impact of mycolactone-BZ combination on immune cell viability, we measured the induction of apoptosis and death in peripheral blood mononuclear cells (PBMCs) subjected to the same drug treatments as patient-derived tumors in Fig. 5. Minimal or no viability loss was observed in T cells, B cells, natural killer cells, dendritic cells and monocytes/macrophages exposed to mycolactone, BZ or both drugs in the conditions tested (**Fig. S5B and Fig. S6**), indicating that the cytotoxicity of the mycolactone-BZ combination is highly selective of MM cells.

### Combining mycolactone with BZ delays MM tumor growth *in vivo*

To evaluate the therapeutic efficacy of the mycolactone-BZ combination *in vivo*, we first analyzed its toxicity in mice. C57BL/6 mice were treated twice a week by intraperitoneal injection of BZ (0.5 mg.Kg-1) alone and/or mycolactone (0.3 mg.Kg-1) during 3 weeks, dosages previously shown to induce anti-MM activity of BZ (24) and anti-inflammatory effects of mycolactone in injected mice (25), respectively. No sign of distress or weight loss could be detected in mice receiving single drugs or the drug combination (data not shown), and their blood cell counts remained unaltered throughout the experiment (**Fig. 6A**), illustrating the safety of these treatment regimen. We next compared the anti-MM activity of mono- and bi-therapies in immunodeficient NOD/SCID/IL2rγ^null^ (NSG) mice. Mice were subcutaneously engrafted with MM.1S cells, and 7 days later they were assigned to four treatment groups receiving (i) 0.5 mg.Kg-1 BZ, (ii) 0.3 mg.Kg-1 mycolactone, (iii) both drugs at these concentrations or (iv) vehicle, twice weekly via the intraperitoneal route. Under these conditions, BZ and mycolactone both induced a minor yet significant delay in tumor growth, compared to vehicle controls (**Fig. 6B**). Notably, the mycolactone-BZ combination was superior to single drug treatments in slowing down MM development. In addition to revealing a therapeutic window for Sec61 blockade in MM, these data confirmed the interest of combining inhibitors of translocon and proteasome in MM therapy.

**Figure 6.**
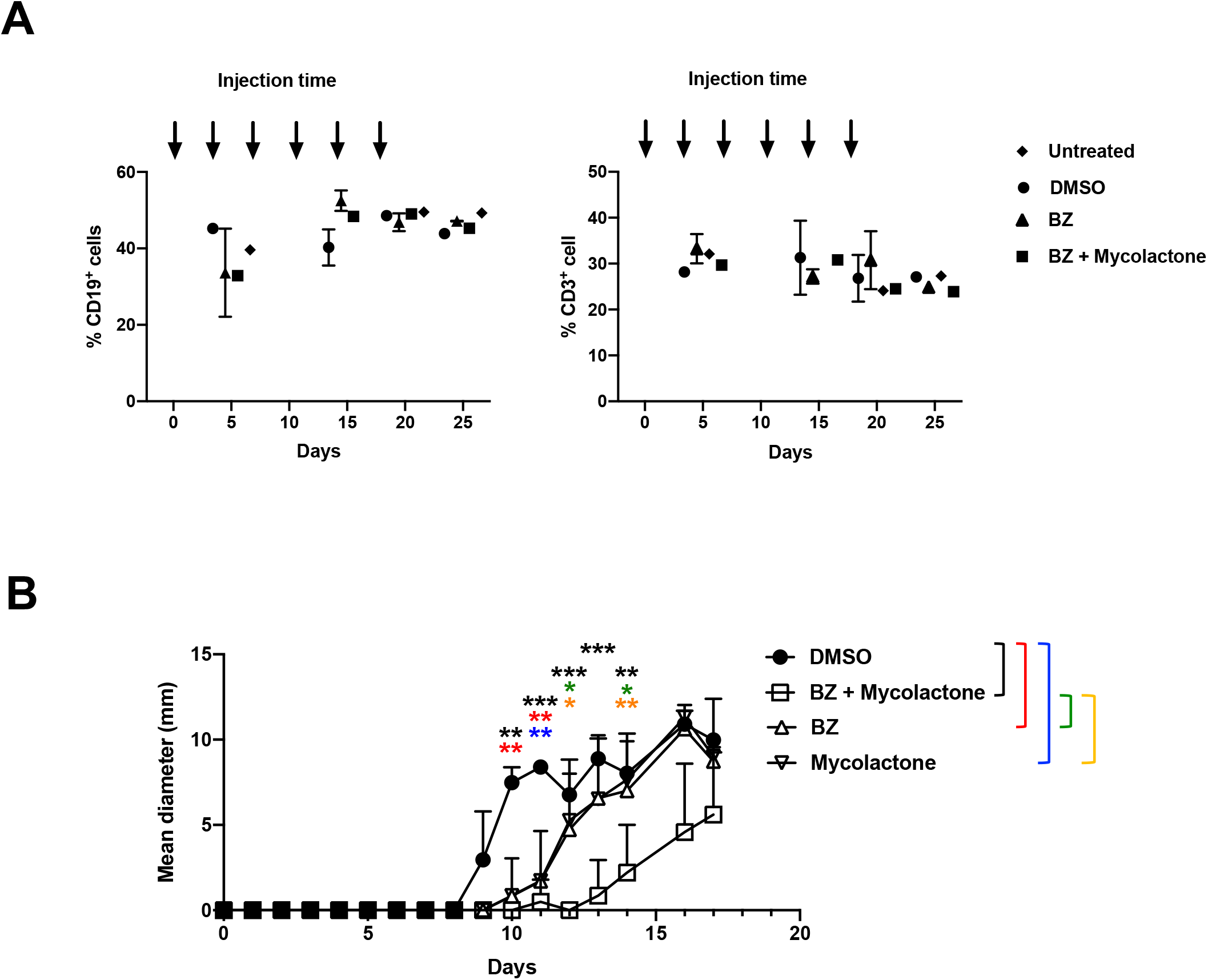
Combining mycolactone with BZ delays MM tumor growth *in vivo*. **(A)** C57BL/6 mice (N = 2 mice/group) were injected intraperitoneally with DMSO, BZ (0.5 mg.Kg^-1^), mycolactone (0.3 mg.Kg^-1^), both drugs at these concentrations or vehicle as controls every 3.5 days. Blood samples were analyzed 5, 15, 20 and 25 days after the first injection and percentages of CD3^+^ and CD19^+^ cells determined by flow cytometry. Results are expressed as % of positive cells among total mononuclear cells, each dot representing the Mean % ± SD of 2 mice. **(B)** NSG mice (N = 9) were injected subcutaneously with 3.10^6^ MM.1S cells at day 0. Seven days later, mice were injected with DMSO, BZ (0.5 mg.Kg^-1^) and/or mycolactone (0.3 mg.Kg^-1^) every 3.5 days intraperitoneally and tumor growth was followed by daily measurement of the tumor diameter. Data are Mean tumor diameters ± SD and represent cumulative data from 3 independent experiments with 3 mice/group. Difference between groups were analyzed with Tukey’s multiple comparison test using Two-way Anova, with mixed effect model for each time points: *\*p<0.05; **p<0.01; ***p<0.001*. Only significant differences are shown.

## Discussion

Although cancer cell line screens have identified Sec61 as a potential target for treatment, the translational potential and therapeutic indications of Sec61 blockers remain largely unknown (26). The first identified Sec61 inhibitor, cotransin, was isolated by screens for molecules decreasing the expression of cell adhesion molecules by a vascular endothelial cell line (27). The high substrate selectivity of cotransin was subsequently used to specifically block cancer cell expression of HER3 (28), a receptor activating oncogenic signaling. Here, we took the opposite approach and sought to exploit the broad, almost total, lack of selectivity of mycolactone for Sec61 substrates (15). Having shown that Sec61 blockade by mycolactone elicits potent ER stress responses (15), we selected the MM cancer model for proof of concept because of its high susceptibility to proteostasis (29). Furthermore, we knew from previous studies that systemically-delivered mycolactone diffuses broadly in injected organisms to accumulate in leukocytes of peripheral blood and lymphoid organs (30, 31), a distribution profile that was adapted to plasma cell targeting. Finally, we reasoned that mycolactone may cause defects in MM cell expression of membrane receptors critically contributing to its dissemination and growth, thereby providing an additional therapeutic benefit *in vivo*. Our *in vitro* investigations using MM cell lines validated these predictions. Moreover, our observations that mycolactone efficiently kills tumors from treatment-naïve and relapsed MM patients *ex vivo* and delays MM xenograft growth in a pre-clinical murine model of disease establish Sec61 as a novel therapeutic vulnerability in MM.

Serum levels of paraproteins, the antibody fragments produced by MM, are used clinically to diagnose MM and monitor its progression (32). In MM cell lines, the level of immunoglobulin light chains released in culture supernatant was dose-dependently decreased by mycolactone after 24 h, and this inhibition closely correlated with the onset of apoptosis at 72 h. This result is in line with our previous demonstration that inhibition of protein secretion on the one hand, and induction of cell apoptosis on the other hand, are early and late effects of mycolactone-mediated Sec61 blockade, respectively (33, 34). It suggests that the prognostic significance of serum paraproteins in patients with MM would not be compromised by administration of Sec61 blockers. Further, mycolactone-mediated inhibition of paraprotein secretion may help limit their pathophysiological effects in treated patients (35).

The introduction of PIs in anti-MM chemotherapies has significantly improved patient survival and PIs now form one of the backbones of treatment (3). It was therefore important to determine whether and how Sec61 blockade interferes with the antitumor activity of PIs. Our data obtained in MM cell lines revealed a synergistic effect between mycolactone and BZ for induction of ER stress and apoptosis, with combination of sub-toxic concentrations of each drug efficiently triggering MM cell death within 24 h *in vitro*. Similar results were obtained with Carfilzomib, indicating that our findings obtained with BZ generalize to second-generation PIs. Mycolactone also synergized with BZ for killing mouse B-ALL and human primary MM cells *ex vivo*, and the mycolactone-BZ combination was superior to single drugs in delaying MM growth in mouse models of disease. Together, these data reveal the potential of Sec61 blockers for combination therapy with PIs.

Today, the gold standard of MM care is a combination of PIs, IMiDs and corticosteroids. While the majority of newly diagnosed patients respond to this 3-drug combination, all eventually develop drug resistance. IMiDs such as lenalidomide operate primarily by targeting Cereblon (CRBN) (36), a substrate receptor of the CRL4 ubiquitin ligase complex. IMiDs binding to CRBN leads to increased proteasomal degradation of pro-survival transcription factors IKZF1/3, thereby promoting MM apoptosis. Notably, low CRBN expression is often associated with IMiD resistance and functional introduction of CRBN mutations in MM cells conferred resistance to lenalidomide *in vitro* (37). In both MM cell lines and primary tumors, mycolactone displayed cytotoxic effects irrespective of the cell basal resistance to BZ. It will be interesting to determine whether mycolactone also kills IMiD-resistant MM, and synergizes with IMiDs for induction of MM cell death.

Mycolactone-PI combinations efficiently killing primary tumor cells displayed minor toxicity to PBMCs from healthy donors and other cell subsets in bone marrow aspirates from MM patients, except for monocytes/macrophages. There is evidence to suggest that in the bone marrow, tumor-associated macrophages operate by supporting MM growth and drug resistance (38). BZ-exposed proinflammatory macrophages were reported to establish a pro-tumorigenic crosstalk with MM cells (39). While additional work will be required to conclude, this strongly suggests that the toxicity of mycolactone-BZ combination in tumor-associated macrophages may actually not be detrimental to MM elimination.

To address the toxicity and resistance issues associated with PI-based treatments, novel approaches using antibody-drug conjugates, bispecific T-cell engager antibodies and Chimeric Antigen Receptor (CAR)-T cells are in test phase. Such strategies target membrane receptors that are specifically expressed by MM cells, such as BCMA (40). Inhibiting Sec61 with cotransin increased the surface expression of BCMA by MM cell lines, resulting in enhanced efficacy a BCMA-targeting antibody-drug conjugate (41). In line with this finding, mycolactone dose-dependently increased BCMA expression in the MM cell lines tested in the present study (Fig. 1). This argues that anti-MM therapies targeting BCMA may benefit from combination with mycolactone.

Here, we used immunodeficient NSG mice to assess the translational potential of mycolactone. Although this setting is widely used to estimate the therapeutic index of drug candidates, it does not allow to evaluate their impact on immune responses. Since Sec61 inhibition strongly affects the functional biology of immune cells (9, 10, 25, 33, 34, 42), it may suppress the development of antitumor immunity. Further investigations using immunocompetent mouse models of MM and B-ALL will be needed to address this hypothesis. Regardless of this limitation, this study reports the first attempt to combine inhibitors of both the proteasome and translocon in Oncology. Our results establish Sec61 blockade as a novel therapeutic approach in MM, synergizing with proteasome inhibition. We used mycolactone for proof of concept, as it displays unprecedented Sec61 inhibition potency and a bodywide distribution *in vivo* (34). However, mycolactone is a complex natural product whose chemical synthesis is a costly, multi-step process making large-scale industrial production challenging (43–45). Structure-activity relationship studies have identified the minimal structural module retaining Sec61 binding activity in mycolactone (9, 25) and the tri-dimensional structure of the Sec6 complex inhibited by mycolactone was recently resolved (46). Our study thus paves the way to structure-based design of drug-like surrogates of mycolactone as novel therapeutics for MM, and potentially other proteostasis-addicted cancers, such as non-small cell lung cancers and certain solid tumors.

## Materials and Methods

### Patients

Patients were seen at Saint-Louis hospital, Paris, France, for a new or known diagnosis of MM. Characteristics of the patients are shown in Table 1. Bone marrow aspirates were performed in the department of Immuno-Hematology of the Saint-Louis Hospital using standard procedures, as part of routine diagnostic work-up. Written informed consent for research was provided by the patients in accordance with the Declaration of Helsinki and French law. All samples containing measurable numbers of viable mononuclear cells were processed and included in data analysis. The protocol received approval of the ethical committee “Groupements hospitaliers universitaires” GHU Nord; N° 23798-2020012709469025 # 121.

### Reagents

Mycolactone was purified from *M. ulcerans* bacterial pellets (strain 1615, ATCC 35840) then quantified by spectrophotometry and stored in ethanol at −20°C protected from light. For *in vivo* experiments, a 4 mM stock was diluted in PBS immediately before injection in animals. For *in vitro* experiments, a 1000 x working solution was prepared by dilution of the ethanol stock in DMSO and stored at −20°C, then thawed and diluted in culture medium immediately before use. BZ purchased from Alfa Aesar (#J60378) was resuspended in DMSO to yield a 10 mM solution and kept at −20°C. A 10 mM solution of Carfilzomib in DMSO was purchased from ApexBio (#A1933) and stored at −20°C. BZ and Carfilzomib were thawed and diluted in culture medium immediately before use. Thapsigargin purchased from Tocris (#1138) was resuspended in DMSO to yield a 10 mM solution and kept at −20°C.

### MM cell line cultures and drug treatment

The MM.1S and KMS-11 cell lines were from J.-C.B. (47). JIM3 cells were kindly provided by Dr. MacLennan (Birmingham Medical School, Birmingham, UK). Cells were cultured in RPMI 1640 medium supplemented with 10% fetal bovine serum (FBS, Dominique Dutcher #S1810-500), 100 Units/mL penicillin + 100 μg/mL streptomycin (Gibco, #15140-122). Exponentially growing MM.1S, JIM3 and KMS-11 cells were plated in 96 well plates at a density of 1.5 10^5^ cells/well (MM-1S) or 10^5^ cells/well (JIM3 and KMS-11). Cells were then treated with mycolactone and/or BZ at the indicated concentrations for 24, 48 or 72 h at 37 °C.

### Lymphomas

Total bone-marrow from 3-5-week-old female C57BL/6 mice was cultured and infected with a retrovirus encoding v-abl kinase to generate immortalized pro-B cell lines as previously described (48, 49). v-abl pro-B cell lines were cultured in RPMI GlutaMAX™ supplemented with 12% heat-inactivated FBS, 10 mM Hepes, and 50 μM 2-mercaptoethanol. Cells were treated with mycolactone and/or BZ as above.

### Flow cytometric analysis of MM cell lines

Following drug treatment, MM cells were pelleted and resuspended in flow cytometry buffer (1× PBS with 1% FBS) then incubated with human FcR blocking reagent (Miltenyi Biotec, # 130-059-901) before staining with diverse fluorescent dye-conjugated monoclonal antibodies: anti-IL-6R/APC (#352806 Biolegend), anti-CD38/BV421 (#303526 Biolegend), anti-CD38/PE (#356604 Biolegend), anti-CD138/APC (#130-117-395 Miltenyi Biotec), anti-CD38/APC-AF750 (#352316 Biolegend), anti-CD40/FITC (#334306 Biolegend), anti-BCMA/PerCp-Cy5.5 (#355710 Biolegend). Cells were stained for 15 min at 4° C, then washed and filtered before data acquisition by Cytoflex flow cytometer (Beckman Coulter). Data were analyzed with the FlowJo software (Tree Star, Ashland, OR). Mean fluorescence intensities (MFI) of isotype controls recommended by the Ab supplier were systematically subtracted to that of each stained sample. Cell viability was assessed by Annexin V exposure and PI incorporation using the FITC-Annexin V/PI kit from Miltenyi Biotec (#130-092-052) following the manufacturer’s protocol. Live cells were characterized as Annexin V^-^ PI^-^, apoptotic cells as Annexin V^+^ PI^-^ and dead cells as Annexin V^+^ PI^+^ and Annexin V^-^ PI^+^ as shown in Figure S1.

### Detection of secreted Immunoglobulin light chain by ELISA

Supernatants from MM cells cultures treated with mycolactone and/or BZ alone were collected after 24 h, and the concentration of free light chain was assessed using the SEBIA FLC lambda ELISA kit (SEBIA, 5101) according to the manufacturer’s recommendations.

### Western blot analyses

Following a 6 h treatment with mycolactone and/or BZ at the indicated concentrations, 2.10^6^ cells were harvested and solubilized at 10^8^ cells/ml for 15 min in ice-cold lysis buffer containing 1% Nonidet P-40, 1% n-dodecyl-{beta}-D-maltoside, 20 mM Tris-HCl, pH 7.5, 150 mM NaCl, 1 mM MgCl2, and 1 mM EGTA in the presence of inhibitors of proteases and phosphatases (10 μg/ml leupeptin, 10 μg/ml aprotinin, 1 mM Pefabloc-sc, 50 mM NaF, 10 mM Na4P2O7, and 1 mM NaVO4). For immunoblot analyses, loadings were normalized on the basis of the total amount of proteins per lane. Proteins were then separated by sodium dodecyl sulfate polyacrylamide gel electrophoresis (SDS-PAGE) using NuPAGE Bis-Tris gels (Thermo Fisher Scientic) and transferred on nitrocellulose membranes (iBlot 2^®^ gel transfer Stacks Nitrocellulose system from Invitrogen). Immune blotting was carried out with the primary antibodies anti-ATF4 (Cell Signaling, #11815), anti GAPDH (Cell Signaling, #2118) overnight at 4°C. After washing, the membranes were incubated with HRP conjugated anti-rabbit and mouse IgG secondary antibodies (Santa Cruz, sc-2004) for 45 min at room temperature. Detection of proteins was performed with the enhanced chemical luminescence (ECL) method using the ECL Prime Western Blotting Reagent and image were acquired on an ImageQuant LAS 4000 Mini (GE Healthcare).

### Gene expression analyses

Following a 6 h treatment with mycolactone and/or BZ at the indicated concentrations, 2.10^6^ cells were harvested and lysed in Trizol (Qiagen). Chloroform was added to the trizol lysates, and the mix was then centrifuged for 15 minutes at 15,000 rcf and 4°C. After centrifugation, the aqueous phase was recovered and mixed with 1.5 volume of ethanol. Total mRNAs were then extracted using RNeasy Plus Mini Kit (Qiagen) according to the manufacturer’s procedure and reverse-transcribed into cDNAs using High Capacity cDNA Reverse Transcription Kit (BD Bioscience) from 1 μg total mRNA according to the manufacturer’s recommendations. The levels of transcription of the mRNAs coding for the genes of interest were assessed using SyberGreen (Power SYBR Green PCR Master Mix, applied biosystem ref: 4367659) and the following primers synthesized by Eurofins genomics: CHOP (forward), GCACCTCCCAGAGCCCTCACTCTCC; CHOP (reverse), GTCTACTCCAAGCCTTCCCCCTGCG; sXBP1 (forward), GGTCTGCTGAGTCCGCAGCAGG; sXBP1 (reverse), GGGCTTGGTATATATGTGG; XBP1total (forward), TGGCCGGGTCTGCTGAGTCCG; XBP1total (reverse), ATCCATGGGGAGATGTTCTGG. Quantitative PCR conditions used were: 50 °C 2 min, 95 °C 10 min, 95 °C 15 s (40 cycles) and 60 °C for 1 minute. The relative quantification was calculated by the 2^-ΔΔCT^ method and the 18S mRNA was used as endogenous control.

### Flow cytometric analysis of MM patient-derived samples and PBMCs

Patient-derived mononuclear cells were isolated from bone marrow aspirate by density gradient centrifugation with Pancoll separating solution (PAN-Biotech, DE). 2.10^5^ cells were then plated in 96 wells plates in culture medium in the absence or the presence of mycolactone or BZ alone or in combination as described in the text for 18 h. Cells were then washed and stained with fluorescent-conjugated anti-CD38 and anti-CD138 antibodies to gate plasma cells, in the presence of Annexin V-FITC and PI for 15 minutes at room temperature. Cells were then acquired using a flow cytometer (Attune NXT, ThermoFisher) immediately after the staining and cell viability was assessed by analysis using FlowJo software (Tree Star, Ashland, OR). PBMCs were isolated from blood samples of healthy donors collected by the French Blood Establishment (EFS), by Pancoll density gradient centrifugation. 2.10^5^ cells were plated in 96 wells plates and treated with mycolactone, BZ, alone or in combination for 18 h. Cells were then washed and stained with anti-CD3/PerCPCy5.5 (#332771 BD Bioscience), anti-CD19/FITC (#302206 Biologend), anti-CD56/APC (#341027 BD Bioscience), anti-CD16/PE-CF594 (#562293 BD Bioscience) and anti-CD11c/PE (#333149 BD Bioscience). Dead cells were identified using LIVE/DEAD Fixable Near IR Dead Cell Stain Kit (#L34975 Invitrogen). Cells were then acquired using a flow cytometer (Cytoflex, Beckman Coulter).

### Mice

Six to eight-week-old female pathogen-free C57BL/6J mice were purchased from Charles River Laboratories. The NOD.Cg-*Prkdc^scid^ Il2rg^tm1Wjl^*/SzJ (NSG, stock number: 005557) mice were purchased from the Jackson laboratory and were used between 6 to 12 weeks of age. All mice were housed at animal facilities of the Institut Pasteur under specific pathogen–free conditions with food and water ad libitum. Mouse experiments were validated by CETEA ethics committee number 0068 (Institut Pasteur, Paris, France, DAP170027) and received the approval of the French Ministry of Higher Education and Research. They were performed in compliance with national guidelines and regulations.

### Mouse experiments

*In vivo* toxicity assays were performed on 8 weeks old C57BL/6J (B6) mice purchased from Charles River laboratories. Mycolactone (0.3 mg.Kg^-1^) and BZ (0.3 mg.Kg^-1^) alone or in combination were diluted in NaCl solution (0.9 % w/v) administrated every 3.5 days via the intraperitoneal route. The toxicity of the treatments was assessed by measuring the percentage of T and B cells in the blood sample collected in tubes containing EDTA (0.05 M) from the tail vein 5, 15, 20 and 25 days after the first injection. Red blood cells were lysed using red blood cells lysing buffer (SIGMA R7757). Leukocytes were stained with anti-mouse CD3/PerCPcy5.5 (#45003182 eBioscience) and CD19/APC (#550992 BD Pharmingen) antibodies and fluorescence data were acquired by Cytoflex flow cytometer (Beckman Coulter). The percentages of T (CD3^+^) and B (CD19^+^) cells were determined using FlowJo software. Experiments assessing the effect of the treatments on the development of xenografted MM tumors were performed in 8 to 12 weeks old NSG mice. 3.10^6^ human MM-1S cells were subcutaneously injected in the right flank of the animals in 200 μL of NaCl solution (0.9 % w/v). Seven days later, mice were injected by intraperitoneal route with mycolactone (0.3 mg.Kg^-1^) and/or BZ (0.3 mg.Kg^-1^), in 200 μL NaCL (0.9 % w/v), or vehicle as control, every 3.5 days. Tumor growth was assessed daily by measuring the size of the tumor with a digital caliper. Tumor size is presented as the average of two perpendicular diameters (millimeters). Mice were sacrificed when the tumor diameter reached 20 mm or whenever the animal shows clinical sign of pain according to ethical guidelines.

### Synergy scores

Synergy between mycolactone and BZ was assessed with the *combenefit* software (21), which calculates scores based on the Loewe additivity model using the dose response of each drug. Loewe synergy score are defined as S_LOEWE_ = Y_obs_-Y_Loewe_, where Y_obs_ is the observed effect of the combination and Y_Loewe_ is the theoretical effect of the combination. Therefore, a S_LOEWE_ > 0, shows that drugs act in synergy. On the contrary, S_LOEWE_ <0, depicts an antagonist effect of the drugs. S_LOEWE_ were plotted as heatmaps and statistical significance analyzed by one-sample Student’s t-test.

### Statistical analyses

Other statistical treatments and graphical representations were performed with the Prism software (v8.4.3, GraphPad, La Jolla, CA) and values of P ≤ 0.05 were considered significant. Detailed information on the statistical test used and number of replicates is provided in figure legends.

## Data availability

The data that support the findings of this study are available within the paper and its supplementary information files.

## Acknowledgements

We would like to thank Jean-David Morel (now at EPFL, Lausanne) for advice regarding the experiments using v-abl cells, and the staff of Animal facility of Institut Pasteur for supervising mouse housing. We also thank Emeline Perthame (Bioinformatics and Biostatistics Hub, Institut Pasteur, USR 3756 IP CNRS) for help with statistical analysis. This work was supported by the Cancéropôle Ile de France (C.D. and J.-C.B; Emergence 2019) and the Fondation Française pour la Recherche contre le Myélome et les Gammapathies (C.D., FFRMG). CD acknowledges core support from Institut Pasteur and INSERM (U1221). L.B. was recipient of a Roux-Cantarini post-doctoral fellowship (Institut Pasteur). A.D. is a BioSPC-Université de Paris PhD student, recipient of a doctoral fellowship from the Ministère français de l’Enseignement Supérieur, de la Recherche et de l’Innovation.

## Author contributions

A.D., C.C., L.B. and V.M. performed the experiments. L.D. provided *v-abl* B cell clones and consultation regarding lymphomas. B.A. and J.-C.B. provided bone marrow aspirates from MM patients and guidance with their handling and flow cytometric analysis. A.D., G.D. and C.D. designed the experiments. Data interpretation and writing of the manuscript were performed by A.D., G.D. and C.D.

## The paper explained

### PROBLEM

Multiple Myeloma is an incurable blood cancer. While the majority of patients initially respond to combination chemotherapies, all eventually develop drug resistance or toxicity and their mean survival rate is only 6 years post diagnosis.

### RESULTS

Human cells interact with their environment via proteins that are either expressed on their surface or secreted into the extracellular environment. The process of delivering these proteins to the membrane wall or outside - the secretion pathway - uses a dedicated distribution network whose gateway is the Sec61 translocon. We recently found that mycolactone, a toxin produced by a bacterial pathogen, operates by blocking Sec61. Using mycolactone we demonstrate in the present study that blocking Sec61 is much more toxic to Multiple Myeloma cells than to normal cells *in vitro* and *in vivo*. Moreover, we show that mycolactone potentiates the activity of drugs forming the backbone of current Multiple Myeloma chemotherapies.

### IMPACT

Our findings identify Sec61 as a therapeutic vulnerability in Multiple Myeloma, and potentially all other cancers needing an active Sec61 translocon to survive. They suggest that treating MM patients with agents blocking Sec61 will augment the efficacy of – and overcome resistance to – current chemotherapies.

## Figure legends

**Figure S1:**
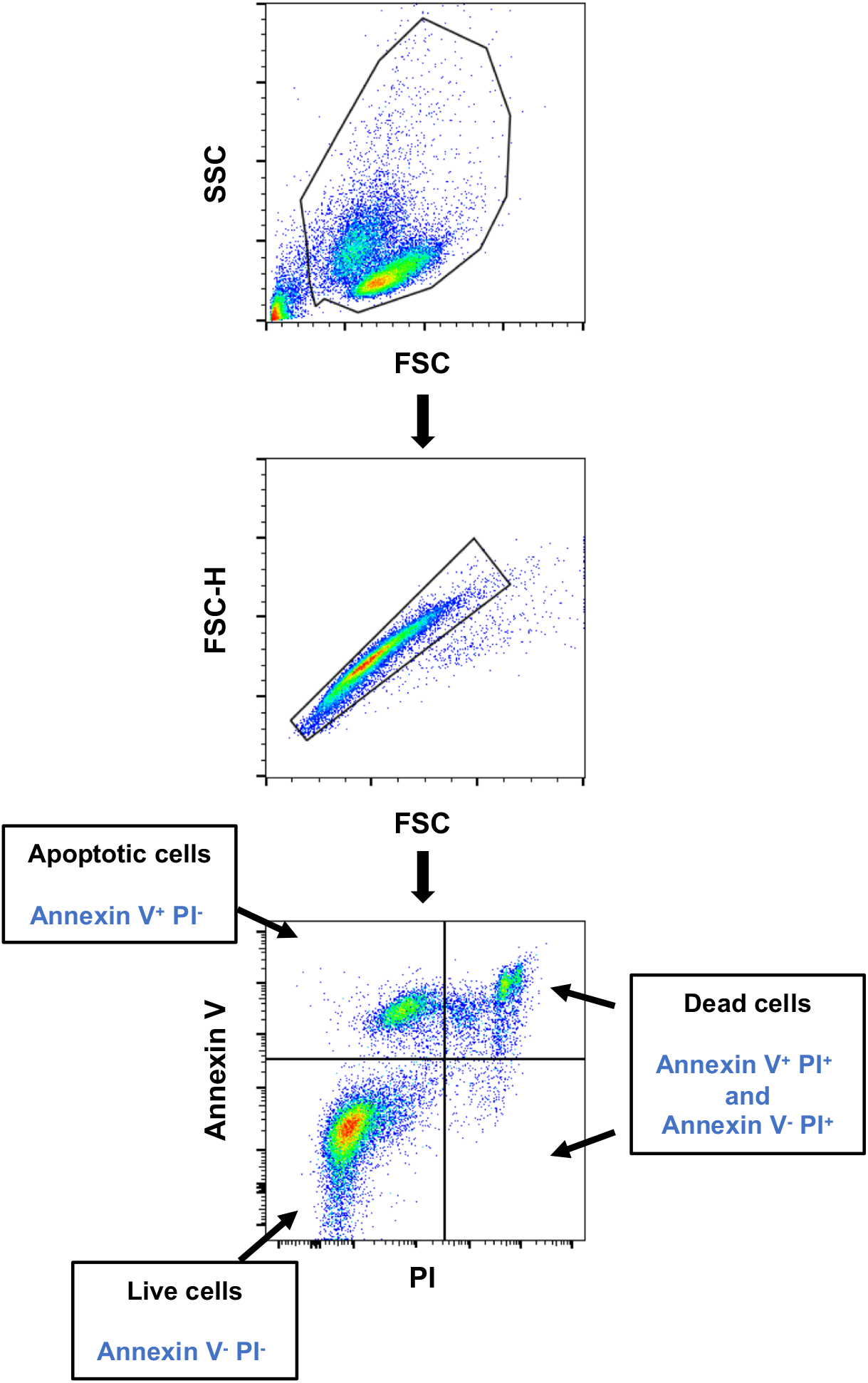
Gating strategy for characterization of live, apoptotic and dead cells. Shown dot plots correspond to MM.1S cell treated with 50 nM mycolactone for 72 h. Live, apoptotic and dead cells were identified by phosphatidylserine exposure (Annexin V staining) and loss of membrane integrity (PI staining), as described in Methods.

**Figure S2:**
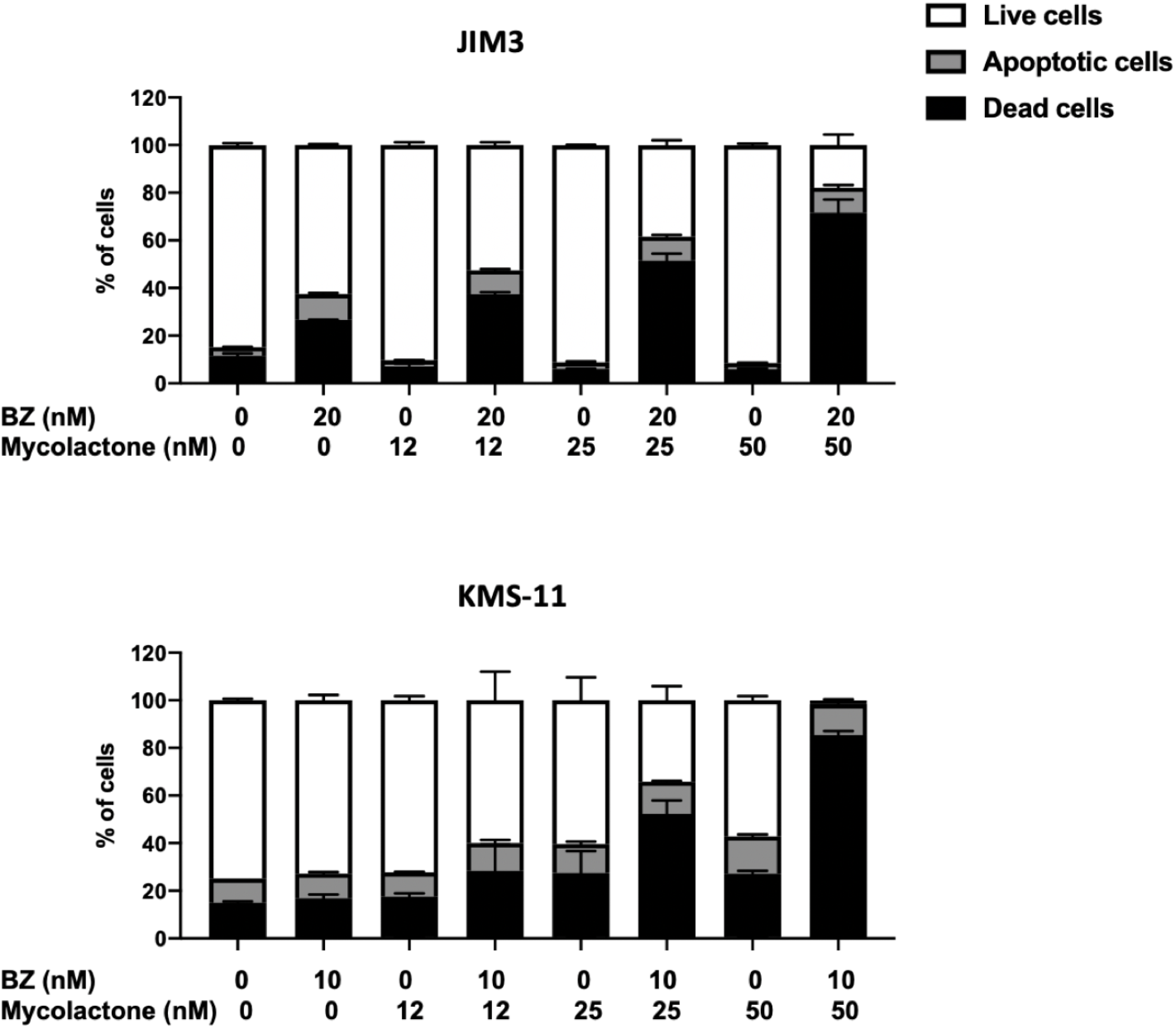
Mycolactone potentiates the effect of BZ in JIM3 and KMS-11. JIM3 and KMS-11 cells were treated with mycolactone and/or BZ at the indicated concentrations for 24 h, and the percentages of live, apoptotic and dead cells were estimated as in Fig. S1. Data are Means ± SD of duplicates, relative to total cell number, and are representative of 2 independent experiments with similar results.

**Figure S3:**
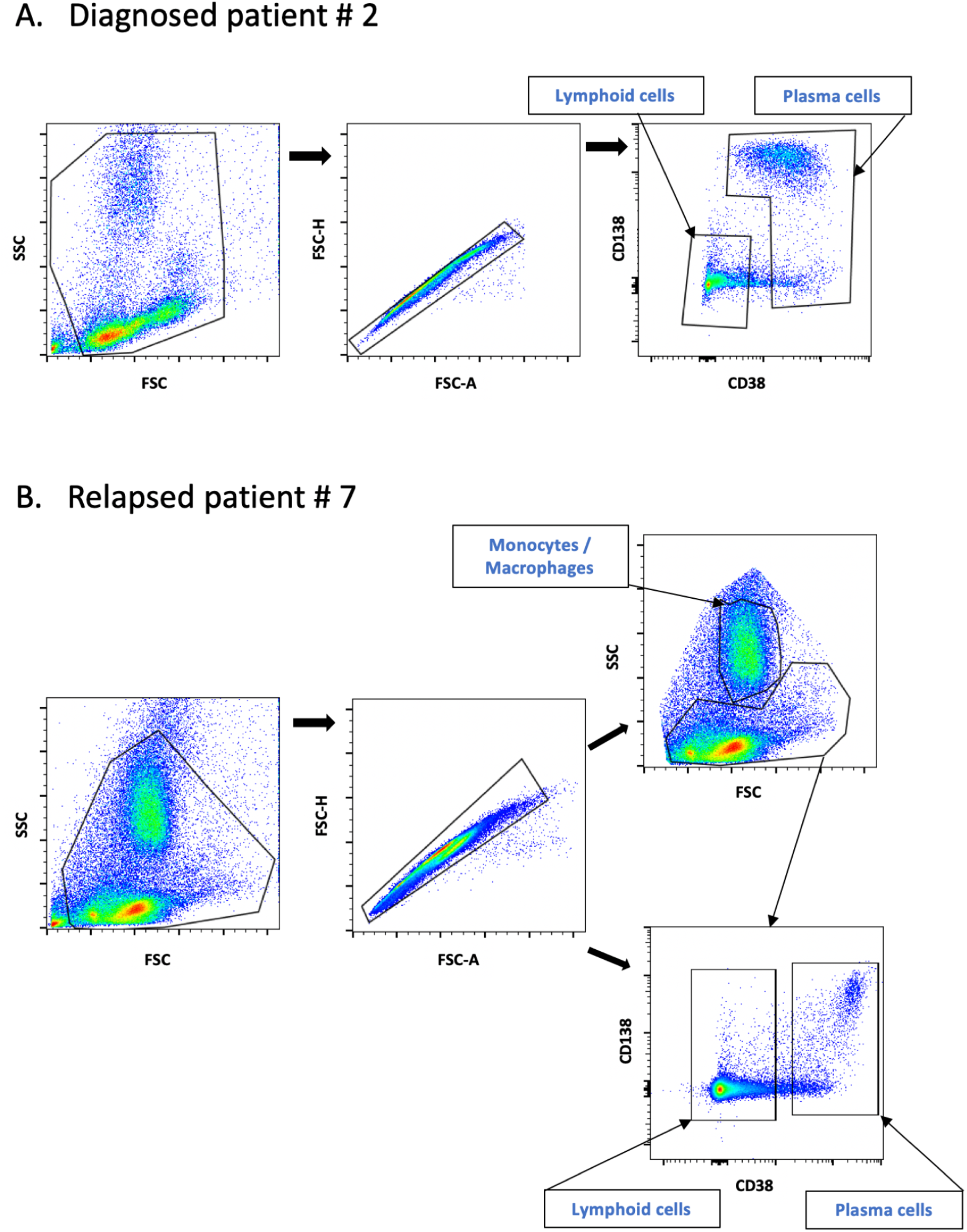
Gating strategy for the identification of MM cells, lymphoid cells and monocytes/macrophages in bone marrow aspirates from MM patients. Diagram illustrating our gating strategy in Pancoll-isolated mononuclear cells from bone marrow aspirates. Shown dot plots correspond to samples from newly diagnosed patient #2 (A) and relapsed patient #7 (B), following staining with anti-CD38 and anti-CD138 antibodies.

**Figure S4:**
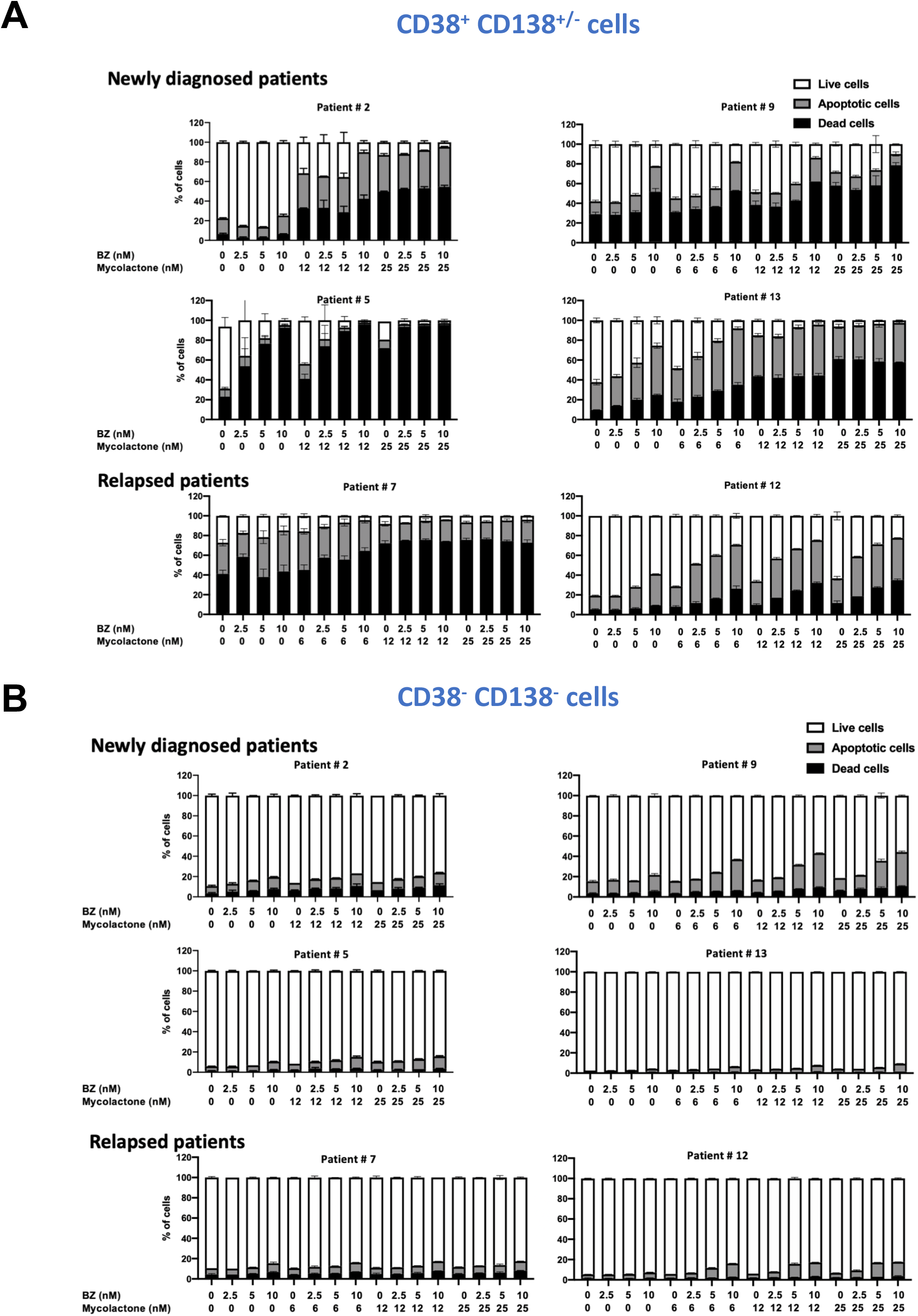
Differential toxicity of mycolactone-BZ combination in MM cells and non-cancerous lymphoid cells in bone marrow aspirates. The percentages of live, apoptotic and dead cells within the MM cell and non-cancerous lymphoid cell subsets after a 18 h treatment with mycolactone and/or BZ are compared in each patient. Mononuclear cells from bone marrow aspirates of newly diagnosed or relapsed MM patients were treated as in Fig. 5. MM cells **(A)** and non-cancerous lymphoid cells **(B)** were identified with the gating strategy depicted in Fig. S3. Data are Mean % ± SD of duplicates, relative to total cell number.

**Figure S5:**
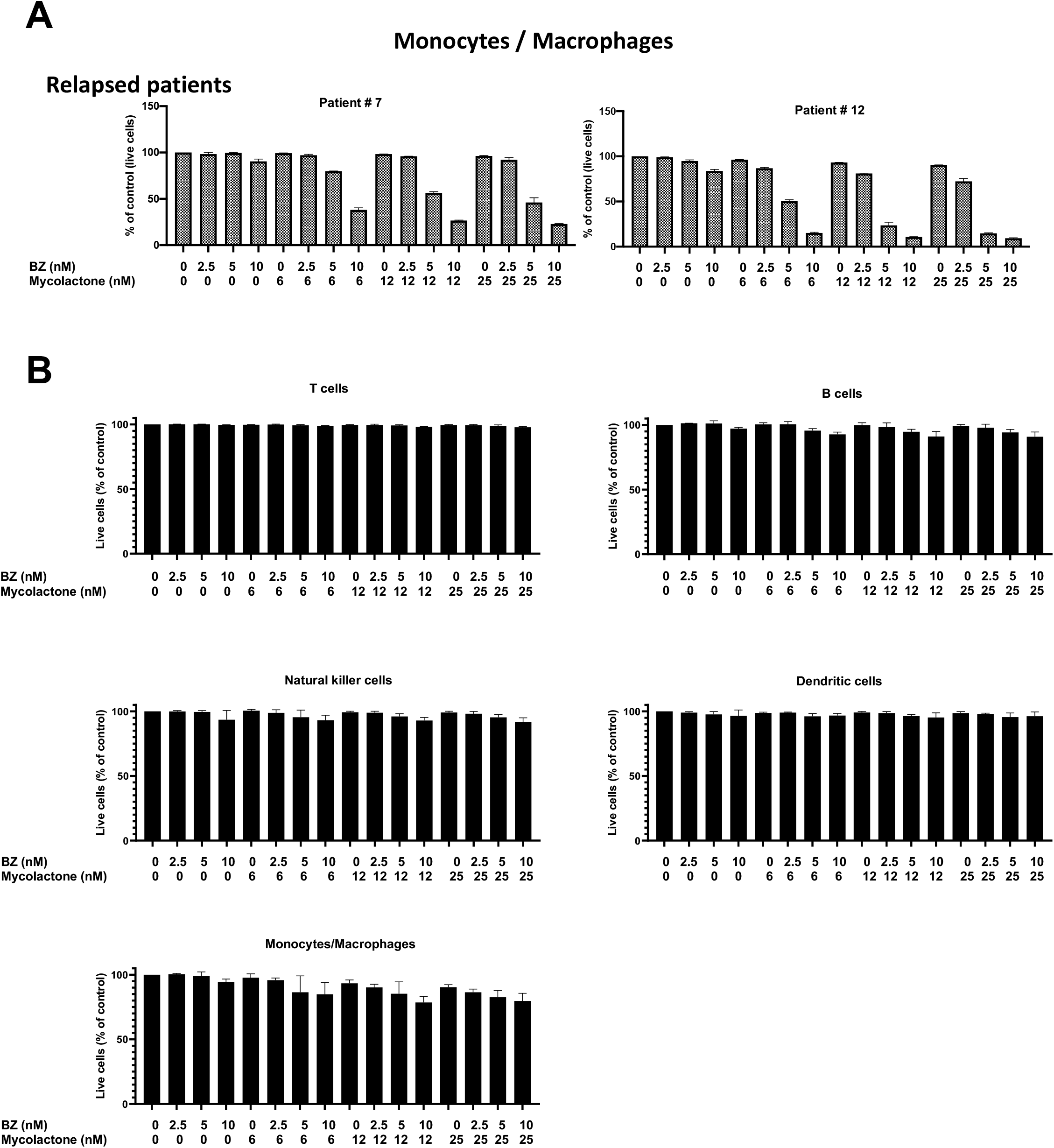
Toxicity of the mycolactone-BZ combination in bone marrow monocytes/macrophages and PBMCs. **(A)** The proportion of live (Annexin V-PI-) cells within the monocyte/macrophage subset of mononuclear cells, after a 18 h treatment with mycolactone and/or BZ is shown for the two relapsed patients. Mononuclear cells from bone marrow aspirates were treated as in Fig. 5. The monocyte/macrophage subset was identified using the gating strategy depicted in Fig. S3. Data are Mean % of live cells +/- SD of duplicates, relative to vehicle-treated controls (Ctrl). **(B)** PBMCs from healthy donors (N = 3) were treated with mycolactone and BZ, alone or in combinations, at the indicated concentration for 18 h. Cells were then labelled with fluorophore conjugated anti-CD3, anti-CD19 and anti-CD11c. T cells, B cells, NK cells dendritic cells and monocytes/macrophages were identified by flow cytometry analysis, using the strategy depicted in Fig. S6. Cell viability was assessed by Annexin V exposure and PI incorporation. Data are pooled measures from three donors and are expressed as Mean percentage of live cells ± SD, relative to untreated cells.

**Figure S6:**
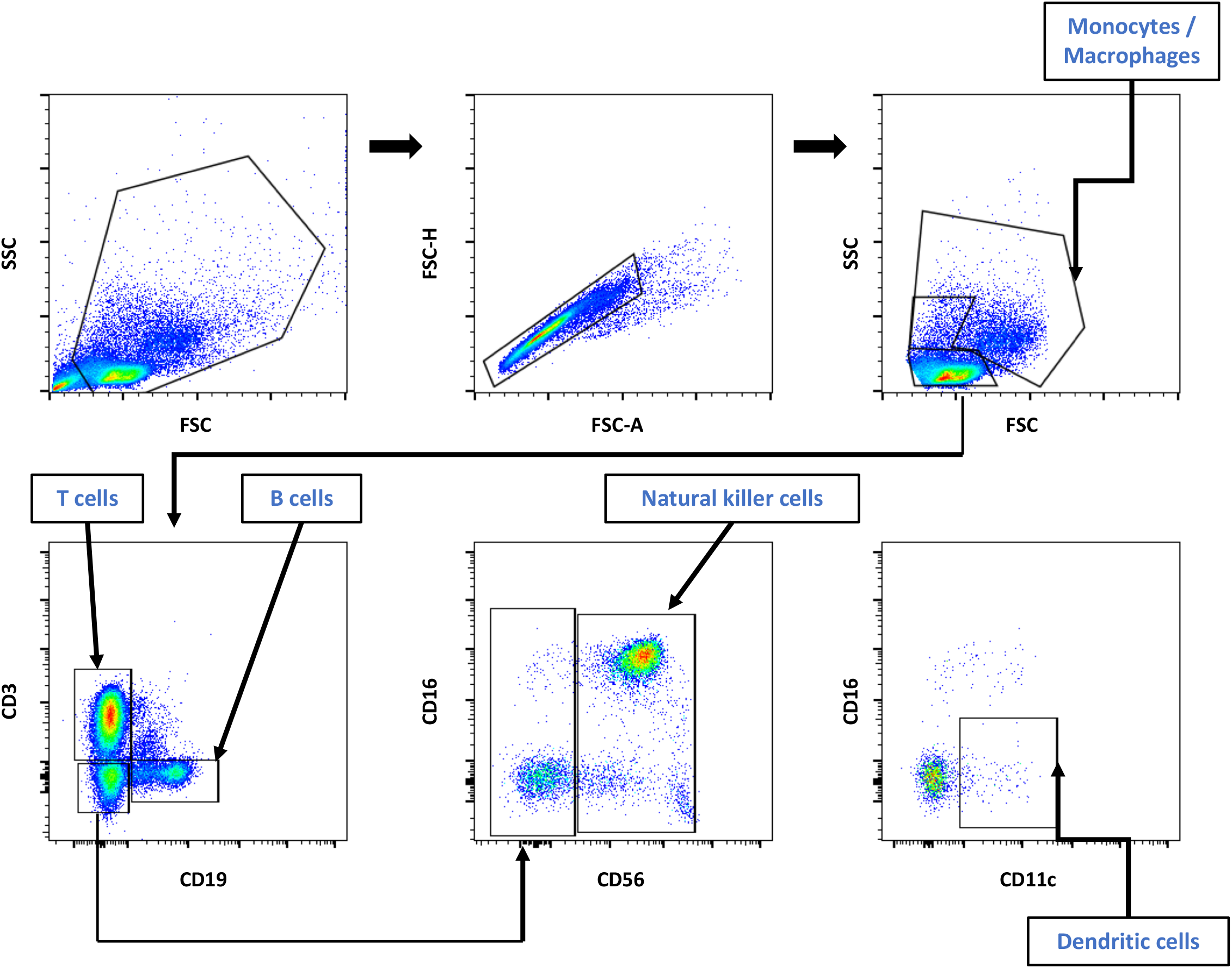
Gating strategy for identification of T cells, B cells, NK cells dendritic cells and monocytes/macrophages in PBMCs. Diagram illustrating our gating strategy in Pancoll-isolated PBMCs from one healthy donor, following staining with anti-CD3, anti-CD19, anti-CD16, anti-CD56 and anti-CD11c antibodies.

